# Sildenafil amplifies calcium influx and insulin secretion in pancreatic β cells

**DOI:** 10.1101/2024.02.17.580848

**Authors:** Naoya Murao, Risa Morikawa, Yusuke Seino, Kenju Shimomura, Yuko Maejima, Yuichiro Yamada, Atsushi Suzuki

## Abstract

Sildenafil, a phosphodiesterase-5 (PDE5) inhibitor, has been shown to improve insulin sensitivity in animal models and prediabetic patients. However, its other metabolic effects remain poorly investigated. This study examines the impact of sildenafil on insulin secretion in MIN6-K8 mouse clonal β cells.

Sildenafil is shown to amplify insulin secretion by enhancing Ca^2+^ influx, an effect that requires other depolarizing stimuli in MIN6-K8 cells but not in K_ATP_ channel-deficient β cells, which are already depolarized. These results indicate that the action of sildenafil is dependent on depolarization and is independent of K_ATP_ channels.

Furthermore, sildenafil-amplified insulin secretion is not inhibited by nifedipine or PDE5 knockdown. Thus, sildenafil stimulates Ca^2+^ influx independently of L-type voltage-dependent Ca^2+^ channels (VDCCs) and PDE5, a mechanism that differs from the known pharmacology of sildenafil and conventional insulin secretory pathways.

Our results reposition sildenafil as an insulinotropic agent that can be used as a potential anti-diabetic medicine or a tool to elucidate the molecular mechanism of insulin secretion.

## 3. INTRODUCTION

Glucose stimulates insulin secretion from pancreatic β cells in a multistep process. First, glucose metabolism increases intracellular ATP levels, which leads to ATP-sensitive K^+^ (K_ATP_) channel closure, resulting in membrane depolarization and the opening of voltage-dependent Ca^2+^ channels (VDCCs). This leads to an increase in intracellular Ca^2+^ (Ca^2+^*i*) and stimulation of insulin exocytosis (Henquin, 2009). Thus, glucose-induced insulin secretion (GIIS) is primarily governed by the opening and closing of K_ATP_ channels in response to glucose metabolism. K_ATP_ channels consist of the pore-forming subunit Kir6.2 encoded by *Kcnj11* and the sulfonylurea receptor SUR1 (Inagaki *et al*., 1995).

In type 2 diabetes, β cells are unable to secrete enough insulin to counteract insulin resistance or excess nutrients, resulting in elevated blood glucose levels. Therefore, enhancement of insulin secretion from β cells is central to the treatment of diabetes. However, the options for clinical insulinotropic agents are currently limited. Of these, sulfonylureas bind to SUR1, which in turn closes the K_ATP_ channels and stimulates insulin secretion. This action is independent of blood glucose levels and, therefore, can cause hypoglycemia as a side effect. There is therefore a need for alternative therapeutic options.

Drug repurposing presents a promising solution for the development of novel insulinotropic medications. Sildenafil is a phosphodiesterase-5 (PDE5) inhibitor primarily used for the treatment of erectile dysfunction and pulmonary arterial hypertension (PAH) (Ghofrani *et al*., 2006). This drug has gained attention as a potential antidiabetic agent owing to its effectiveness in preclinical and clinical studies. Sildenafil has been reported to enhance insulin sensitivity in high-fat diet-fed mice (Ayala *et al*., 2007) and prediabetic patients (Ramirez *et al*., 2015). Additionally, sildenafil promotes vascular relaxation in diabetic rats (Schäfer *et al*., 2009) and is considered beneficial for treating vascular dysfunction in diabetic patients (Zimmermann *et al*., 2020).

However, the effect of sildenafil on insulin secretion from pancreatic β cells remains unclear. This study investigates the effect of sildenafil on insulin secretion using MIN6-K8, a mouse clonal β cell line, as a model. Our results indicate that sildenafil promotes insulin secretion by increasing calcium influx and that this effect is not mediated by PDE5 or L-type VDCCs. This suggests a novel mechanism of action for sildenafil in β cells that differs from its known pharmacology.

## 4. MATERIALS AND METHODS

### 4.1 Cell lines

MIN6-K8 cells were established as previously described (Iwasaki *et al*., 2010) and were kindly provided by Professor Junichi Miyazaki (Osaka University). *Kcnj11*^*-/-*^ β cells (clone *Kcnj11*^-/-^ βCL1) are β cells deficient in Kir6.2, established by sub-cloning MIN6-K8 cells transfected with Cas9 nickase and guide RNA pairs targeting mouse *Kcnj11*, as described previously (Oduori *et al*., 2020). All cells were cultured in Dulbecco’s modified Eagle’s medium (DMEM) containing 4500 mg/L glucose (Sigma-Aldrich, St. Louis, MO, USA, Cat# D5796) supplemented with 10% fetal bovine serum (FBS) (BioWest, Nuaillé, France, Cat# S1400-500) and 5 ppm 2-mercaptoethanol. The cells were maintained at 37 °C with 5% CO2.

### 4.2 Reagents

Krebs-Ringer bicarbonate buffer-HEPES (133.4 mM NaCl, 4.7 mM KCl, 1.2 mM KH2PO4, 1.2 mM MgSO4, 2.5 mM CaCl2, 5 mM NaHCO3, 10 mM HEPES) containing 0.1% bovine-serum albumin (Sigma-Aldrich, St. Louis, MO, USA, Cat# A6003) and 2.8 mM glucose (2.8G-KRBH) adjusted to pH 7.4 was used in insulin secretion and Ca^2+^ imaging experiments. Additional glucose (final concentration, 11.1 mM), sildenafil (Tokyo Chemical Industry, Tokyo, Japan, Cat# S0986), and glimepiride (Tokyo Chemical Industry, Tokyo, Japan, Cat# G0395) were added to KRBH during the stimulation period. Nifedipine (FUJIFILM Wako Pure Chemical, Osaka, Japan, Cat# 14505781), diazoxide (Tokyo Chemical Industry, Tokyo, Japan, Cat# D5402), and thapsigargin (FUJIFILM Wako Pure Chemical, Osaka, Japan, Cat# 209-17281) were added during the pre-incubation and stimulation periods. The reagents used for stimulation were stored as a 1000× concentrate in dimethyl sulfoxide (DMSO) (FUJIFILM Wako Pure Chemical, Osaka, Japan, Cat# 041-29351) and diluted with KRBH shortly before the experiment. An equal volume of DMSO was added to the vehicle control. Ca^2+^-free KRB was formulated by replacing CaCl2 with an equivalent concentration of MgCl2 and adding 0.2 mM EGTA (NACALAI TESQUE, Kyoto, Japan, Cat# 15214-21).

### 4.3 Insulin secretion

Insulin secretion was measured using the static incubation method as described previously (Murao *et al*., 2022) with slight modifications. Briefly, cells were seeded in 24-well plates at a density of 5 × 10^5^ cells/well and cultured for 48 h. On the day of measurement, the cells were subjected to three successive washes with 2.8G-KRBH, followed by a pre-incubation period of 30 min with 300 μL/well of 2.8G-KRBH. Subsequently, the supernatant was replaced with 300 μL/well of fresh KRBH containing the specified stimulations and incubated for 30 min at 37 °C.

The reaction was terminated by cooling the plate on ice for ten minutes, after which the entire supernatant was collected for the quantification of released insulin using the homogeneous time-resolved fluorescence assay (HTRF) Insulin Ultrasensitive kit (Revvity, Waltham, MA, USA, Cat# 62IN2PEH) in accordance with the manufacturer’s instructions. Fluorescence was measured using an Infinite F Nano+ microplate reader (Tecan, Zürich, Switzerland).

### 4.4 Imaging of intracellular Ca^2+^

Cells were seeded in a 35 mm glass-bottom dish (Matsunami Glass, Osaka, Japan, Cat# D11530H) at a density of 1.28 × 10^5^ cells/dish and cultured for 48 h. Subsequently, the cells were loaded with 1 μM Fluo-4 AM (Dojindo, Kumamoto, Japan, Cat# F312) in 2.8G-KRBH for 20 min at 37 °C in room air. Following a brief washing, cells were loaded with 1 mL of fresh 2.8G-KRBH and basal recordings were performed for 300 s (from time -300 to 0). Immediately after the addition of 1 mL KRBH supplemented with stimulations at 2× concentration, recordings were resumed for another 600 s (from time 0 to 600) with a time interval of 2 s.

Time-lapse images were obtained using a Zeiss LSM 980 Airyscan2 inverted confocal laser scanning super-resolution microscope equipped with a Plan Apo 40×, 1.4 Oil DICII objective lens (Carl Zeiss Microscopy, Jena, Germany). The cells were excited at 488 nm laser with 0.3 % output power, and fluorescence emission was measured at 508-579 nm. During observation, the cells were maintained at 37 °C using an incubator XLmulti S2 DARK (Pecon, Erbach, Germany).

Images were acquired in the frame mode at a rate of 2 frames per second and with an image size of 212.2 × 212.2 μm (512 × 512 pixels). The obtained images were analyzed using the ZEN 3.0 imaging software (Carl Zeiss Microscopy, Jena, Germany, RRID:SCR_021725). Cells were randomly chosen for analysis for each stimulation, and the number of cells analyzed is indicated in the figure legends. The fluorescence intensity of the entire cell body (F) was monitored and normalized to the average fluorescence intensity between -300 and 0 s (F0). The amplitude of Ca^2+^ responses was quantified as the incremental area under the curve (iAUC) using F/F0 = 1 as the baseline.

### 4.5 Knockdown of *Pde5a* using small interfering RNA (siRNA)

siRNAs targeting *Pde5a* (Dharmacon, Lafayette, CO, USA, Cat# M-041115-00-0005) and non-targeting siRNA (Dharmacon, Lafayette, CO, USA, Cat# D-001206-14-50) were reverse-transfected using the DharmaFECT 2 transfection reagent (Dharmacon, Lafayette, CO, USA, Cat# T-2002-03). Briefly, a complex of siRNA and DharmaFECT 2 was prepared in serum-free DMEM (Sigma-Aldrich, St. Louis, MO, USA, Cat# D5796) at a volume of 100 μL/well **according** to the manufacturer’s instructions. Cells were resuspended in complete culture media at 1.25 × 10^6^ cells/mL. The cell suspension was then combined with siRNA/DharmaFECT 2 complex and seeded in 24-well plates at 5 × 10^5^ cells/500 μL/well. The final concentrations of siRNA and DharmaFECT 2 were 40 nM and 0.4%, respectively. Insulin secretion or RT-qPCR experiments were performed after a 48-hour culture.

### 4.6 RT-qPCR

cDNA was prepared from 48-hour cultured cells using CellAmp Direct Lysis and RT set (Takara Bio, Shiga, Japan, Cat# 3737S/A) according to the manufacturer’s instructions. Quantitative real-time PCR was performed on a QuantStudio 7 Flex system (Thermo Fisher Scientific, Waltham, MA, USA, RRID:SCR_020245) using TaqMan Universal Master Mix II with UNG (Thermo Fisher Scientific, Waltham, MA, USA, Cat# 4440038) and Taqman probes: *Pde5a* (Cat# Mm00463177_m1) and *Tbp* (Cat# Mm01277042_m1). Relative gene expression of *Pde5a* was calculated using the 2^-ΔΔCT^ method and normalized to *Tbp*.

### 4.7 Statistical Analysis

Sample sizes were estimated from the expected effect size based on previous experiments. No randomization or blinding was used. For insulin secretion and RT-qPCR experiments, *n* represents the number of biological replicates of cells grown in individual wells. For Ca^2+^ measurements, *n* represents the number of different single cells analyzed. Data are shown as the mean ± standard error of the mean (SEM) along with the plot of individual data points. For statistical comparisons between two groups, a two-tailed unpaired Welch’s unpaired *t*-test was used. For more than three groups, one-way analysis of variance (ANOVA) was followed by pairwise comparisons corrected using Dunnett’s method. P-values smaller than 0.05 were considered statistically significant and are indicated in the figures. P-values greater than 0.05 **are** indicated in the figures. The statistical analyses used are indicated in the figure legends. No statistical methods were used to determine whether the data met the assumptions of the statistical approach. Statistical analyses were performed using GraphPad Prism 9 (Graphpad Software, Boston, MA, USA, https://www.graphpad.com; RRID:SCR_002798).

## 5. RESULTS

### 5.1 Sildenafil amplifies insulin secretion from β cell lines

The effect of sildenafil on insulin secretion was investigated using MIN6-K8 cells. Sildenafil at concentrations greater than 10 μM dose-dependently enhanced insulin secretion at stimulatory levels (11.1 mM) of glucose (Figure 1A). Sildenafil displayed no effect at basal levels (2.8 mM) of glucose (Figure 1B), indicating that the insulinotropic effect of sildenafil is glucose-dependent.

**Figure 1.**
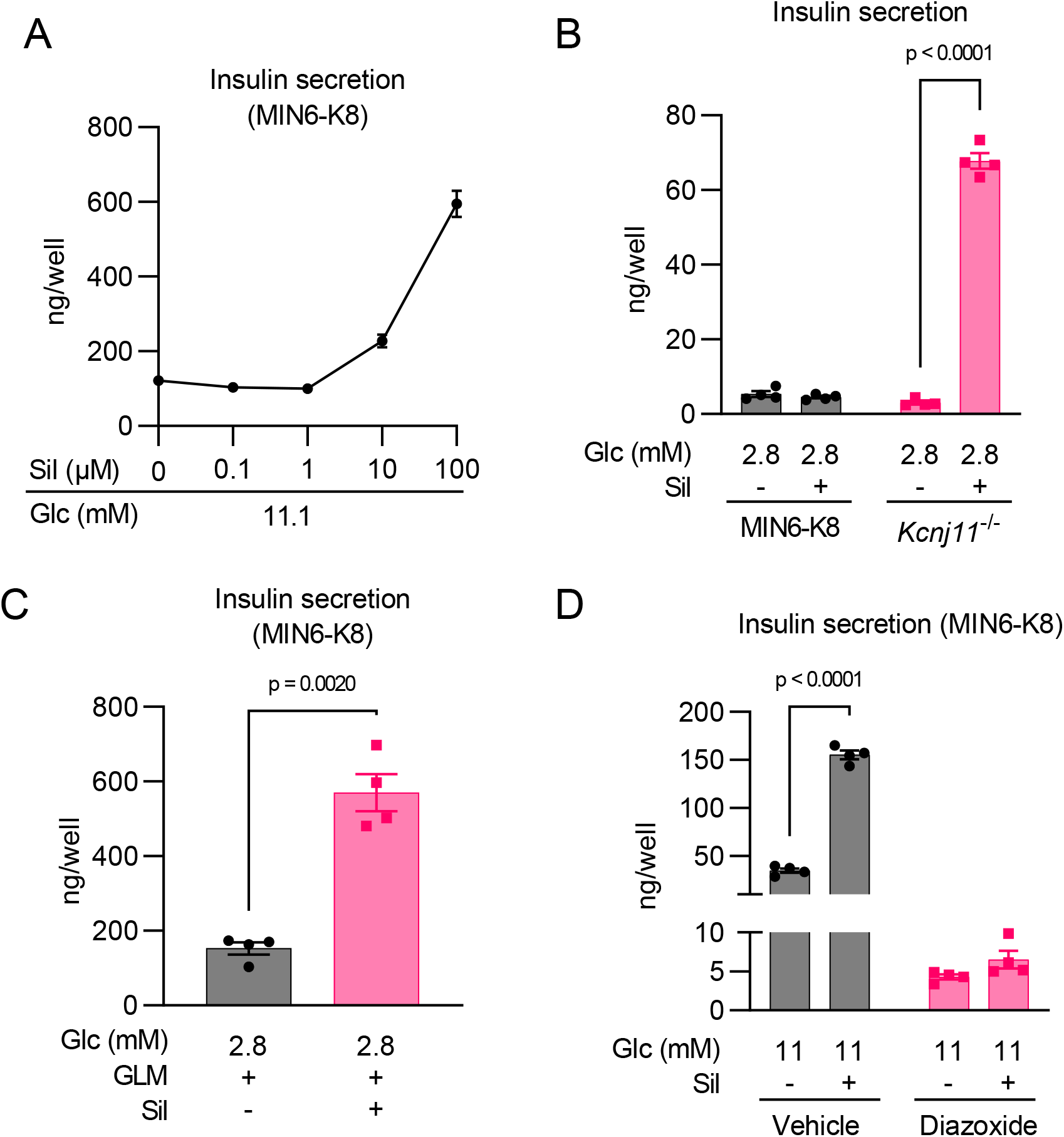
Sildenafil amplifies insulin secretion from β cell lines in a depolarization-dependent manner. (A) Dose-dependent effects of sildenafil on insulin secretion at 11.1 mM glucose in MIN6-K8 cells n = 4. (B) Effects of sildenafil on insulin secretion at 2.8 mM glucose in MIN6-K8 and *Kcnj11*^-/-^ β cells. n = 4. (C) Effect of sildenafil on insulin secretion in the presence of glimepiride at 2.8 mM glucose in MIN6-K8 cells. n = 4. (D) Effect of diazoxide on sildenafil-amplified insulin secretion in MIN6-K8 cells. n = 4. Data are presented as the mean ± standard error of the mean (SEM). 2.8G, 2.8 mM glucose; 11.1G, 11.1 mM glucose. The reagents were added to achieve the following final concentrations unless otherwise specified: sildenafil (Sil) - 100 μM, glimepiride (GLM) - 1 μM, and diazoxide – 100 μM. Statistical comparisons were performed using Welch’s unpaired two-tailed t-test for (B), (C), and (D).

To assess whether sildenafil enhances insulin secretion by facilitating K_ATP_ channel closure, its efficacy was tested in K_ATP_ channel-deficient β cells. We previously generated *Kcnj11*^-/-^ β cells, in which K_ATP_ channel activity is absent and the cell membrane is depolarized continuously regardless of extracellular glucose levels (Oduori *et al*., 2020). Sildenafil significantly increased insulin secretion in *Kcnj11*^-/-^ β cells even at 2.8 mM glucose (Figure 1B), demonstrating that sildenafil-amplified insulin secretion is independent of K_ATP_ channel activity.

Similarly, in MIN6-K8 cells, sildenafil amplified insulin secretion at 2.8 mM glucose in the presence of the sulfonylurea glimepiride, which inhibits K_ATP_ channel activity (Figure 1C). In contrast, sildenafil’s insulinotropic effect at 11 mM glucose was lost in the presence of diazoxide (Figure 1D), a K_ATP_ channel opener that hyperpolarizes β cells (Gribble and Reimann, 2003; Rorsman and Ashcroft, 2018). These observations suggest that sildenafil is effective only when the plasma membrane is depolarized by other factors such as high glucose, sulfonylureas, and *Kcnj11* knockout.

### 5.2 Sildenafil potentiates influx of extracellular Ca^2+^

We then investigated the association between sildenafil-amplified insulin secretion and Ca^2+^*i* using Fluo-4 imaging.

In MIN6-K8 cells, sildenafil augmented the increase in Ca^2+^*i* induced by 11.1 mM glucose but had no apparent effect at 2.8 mM glucose (Figures 2A-B). In *Kcnj11*^-/-^ β cells, sildenafil increased Ca^2+^*i* levels at 2.8 mM glucose (Figures 2C-D). These trends accord with the pattern of insulin secretion shown in Figure 1, suggesting that sildenafil-amplified insulin secretion involves augmented Ca^2+^*i* responses.

**Figure 2.**
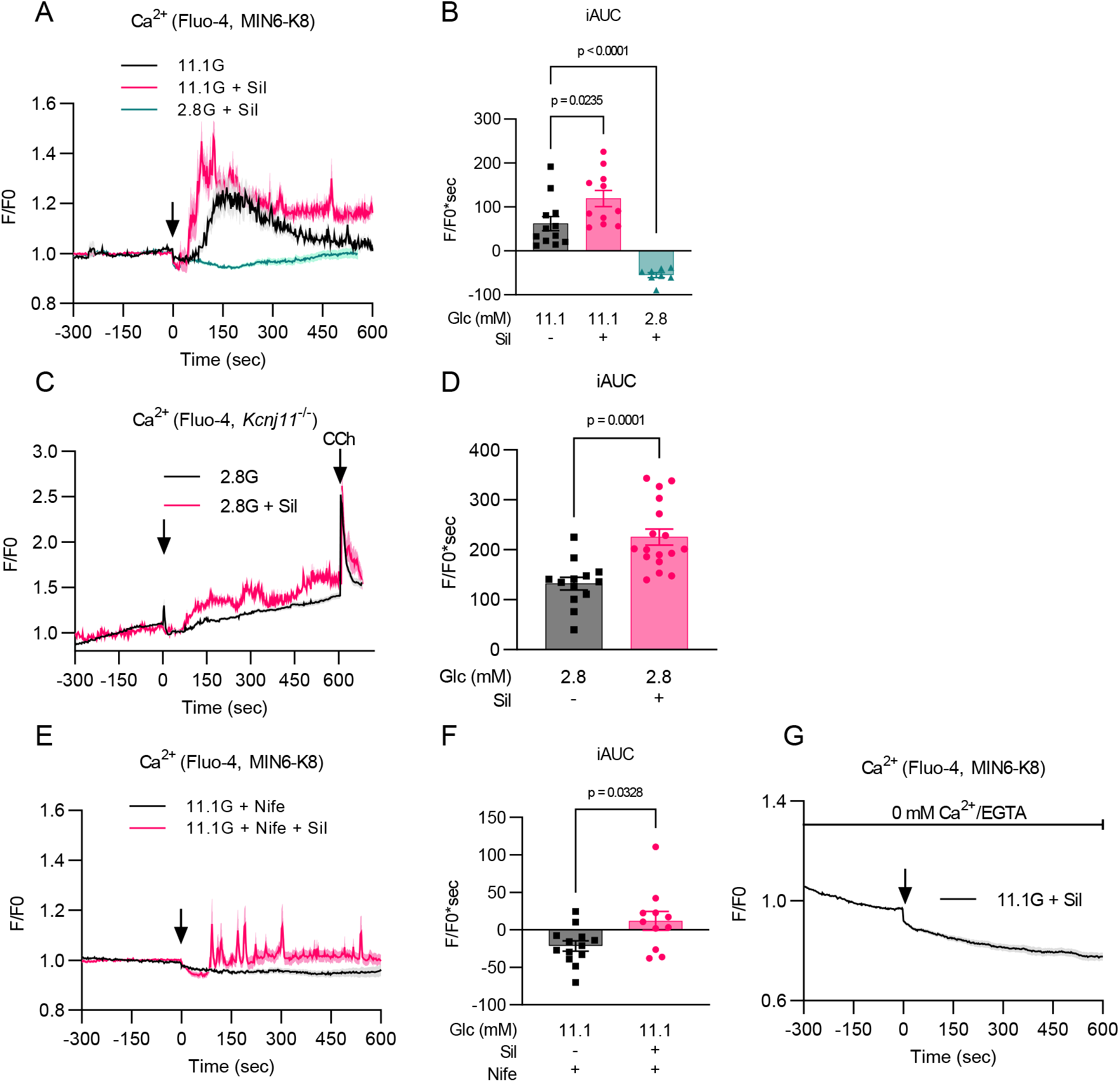
Sildenafil potentiates the influx of extracellular Ca^2+^. Intracellular Ca^2+^ levels were measured using Fluo-4. The time course of normalized fluorescence intensity at 508-579 nm is indicated in (A), (C), (E), and (G). The black arrow indicates the addition of the indicated stimulations at time = 0. The magnitude of Ca^2+^ responses was quantified as iAUC in (B), (D), and (F). (A) (B) Effect of sildenafil on intracellular Ca^2+^ in MIN6-K8 cells. 2.8G + Sil: n = 8; 11.1G: n = 12; 11.1G + Sil: n = 11. (C) (D) Effect of sildenafil on intracellular Ca^2+^ in *Kcnj11*^-/-^ β cells. Carbachol was added at time = 600 as positive control. 11.1G + Nife: n = 13; 11.1G + Nife + Sil: n = 11. (E) (F) Effect of nifedipine on sildenafil-induced Ca^2+^ response in MIN6-K8 cells. 11.1G + Nife: n = 13; 11.1G + Nife + Sil: n = 11. (G) Effect of sildenafil on intracellular Ca^2+^ under extracellular Ca^2+^-free conditions in MIN6-K8 cells. 2.8G + Sil: n = 8; 11.1G: n = 12; 11.1G + Sil: n = 11. Data are presented as the mean ± SEM. The SEM is indicated by shaded regions (A), (C), (E), and (G), as well as by error bars elsewhere. 2.8G, 2.8 mM glucose; 11.1G, 11.1 mM glucose. The reagents were added to achieve the following final concentrations: sildenafil (Sil) - 100 μM, nifedipine (Nife) - 10 μM, carbachol (CCh) - 50 μM, and EGTA - 0.2 mM. Statistical comparisons were made using one-way ANOVA with Dunnett’s post hoc test in (B), and Welch’s unpaired two-tailed t-test for (D) and (F).

Nifedipine, an inhibitor of L-type VDCCs, eliminated the response to 11.1 mM glucose but did not completely inhibit the response to a combination of sildenafil and 11.1 mM glucose (Figures 2E-F). This combination maintained a higher baseline and produced multiple spikes in Ca^2+^ traces even in the presence of nifedipine (Figure 2E). These responses were abolished upon removal of Ca^2+^ from the stimulation buffer (Figure 2G). Consistently, nifedipine treatment substantially lowered insulin secretion by 11.1 mM glucose compared to vehicle alone but did not block the responsiveness to sildenafil (Figure 3A), which was increased as expressed by fold change (Figure 3B). In contrast, Ca^2+^-free buffer abrogated sildenafil responsiveness (Figure 3A-B). Thus, sildenafil-induced Ca^2+^*i* response and insulin secretion are dependent on extracellular Ca^2+^ but not on L-type VDCCs.

**Figure 3.**
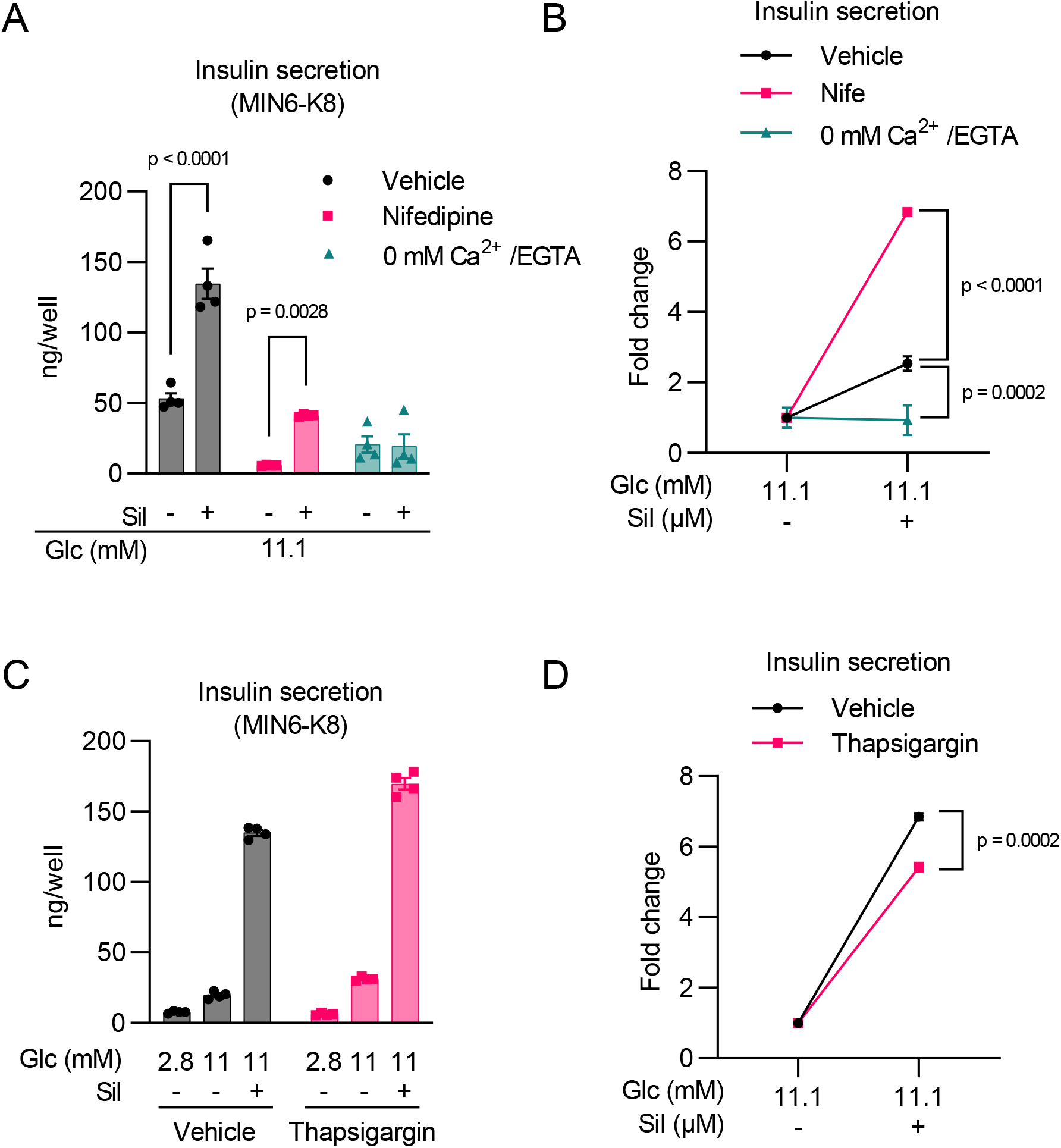
Sildenafil-amplified insulin secretion is dependent on extracellular Ca^2+^. (A) (B) Effects of nifedipine or extracellular Ca^2+^-free conditions on sildenafil-amplified insulin secretion in MIN6-K8 cells. n = 4. The data is presented in its original value in (A) and as fold change over 11.1 mM glucose in (B). (C) (D) Effect of thapsigargin on sildenafil-amplified insulin secretion in MIN6-K8 cells. n = 4. The data is presented in its original value in (C) and as fold change over 11.1 mM glucose in (D). Data are presented as the mean ± SEM. The reagents were added to achieve the following final concentrations: sildenafil (Sil) - 100 μM, nifedipine (Nife) - 10 μM, EGTA - 0.2 mM, and thapsigargin - 1 μM. Statistical comparisons were performed using one-way ANOVA with Dunnett’s post-hoc test for (B) and Welch’s unpaired two-tailed t-test for (A) and (D).

We also assessed the role of intracellular Ca^2+^ stores using thapsigargin, an inhibitor of sarcoplasmic/endoplasmic reticulum Ca^2+^-ATPase (SERCA), to deplete intracellular Ca^2+^ *i* stores. Thapsigargin only marginally affected insulin secretion, with a slight decrease in sildenafil responsiveness as measured by the fold change (Figures 3C-D). This result indicates that intracellular Ca^2+^ stores are dispensable for the sildenafil-induced Ca^2+^ response.

### 5.3 Sildenafil-amplified insulin secretion is independent of PDE5

We then investigated whether PDE5, the original molecular target of sildenafil, plays a role in its insulinotropic effect. Using siRNA to knock down *Pde5a*, we successfully decreased its transcript levels by approximately 50% (Figure 4A). However, this knockdown had no inhibitory effect on sildenafil-enhanced insulin secretion and actually appeared to increase sildenafil responsiveness (Figures 4B-C). These findings indicate that the insulinotropic effect of sildenafil is independent of PDE5 inhibition.

**Figure 4.**
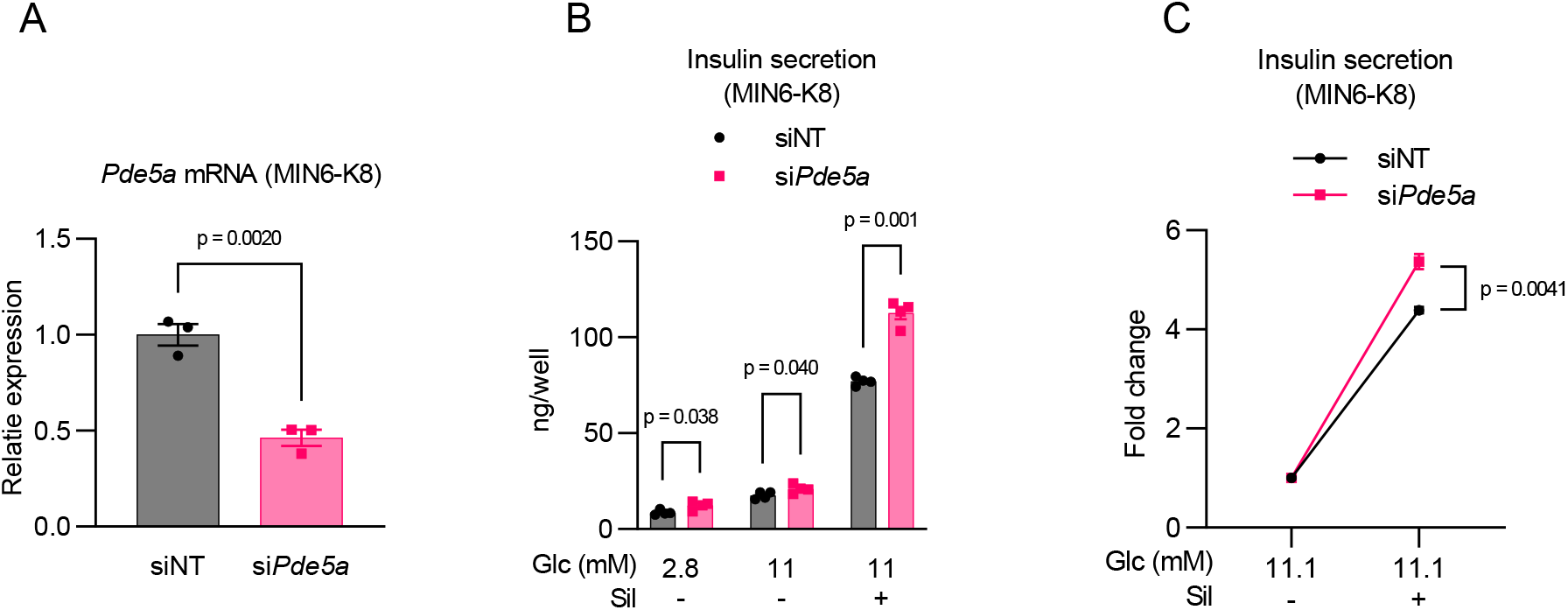
PDE5 is not involved in sildenafil-amplified insulin secretion. (A)Knockdown efficiency of *Pde5a* was assessed by RT-qPCR. mRNA levels were normalized to siNT (non-targeting siRNA)-treated cells. n = 3. (B) (C) Effect of *Pde5a* knockdown on sildenafil-amplified insulin secretion. n = 4. The data is presented in its original value in (B) and as fold change over 11.1 mM glucose in (C). Data are presented as the mean ± SEM. Sildenafil (Sil): 100 μM. Statistical comparisons were made using Welch’s unpaired two-tailed t-test.

## 6. DISCUSSION

In the present study, we show that (1) sildenafil enhances insulin secretion by augmenting extracellular Ca^2+^ influx independently of L-type VDCCs; (2) furthermore, sildenafil-amplified insulin is dependent on depolarization but independent of K_ATP_ channels; and that (3) sildenafil-amplified insulin secretion is not mediated by the PDE5-inhibitory effects of sildenafil.

This is the first report of a direct effect of sildenafil on pancreatic β-cells and suggests its favorable therapeutic properties as an insulinotropic agent in the treatment of diabetes. Research has suggested that K_ATP_ channel activity is impaired in diabetic β cells (Oduori *et al*., 2020; Nichols *et al*., 2022). Unlike sulfonylureas and glinides, which are ineffective in K_ATP_ channel-inactivated β cells, sildenafil remains effective. In addition, the insulinotropic effect of sildenafil is dependent on glucose levels, thereby reducing the likelihood of hypoglycemia as a potential side effect. Moreover, the beneficial effects of sildenafil on insulin sensitivity (Ayala *et al*., 2007; Ramirez *et al*., 2015) and vascular function (Schäfer, *et al*. 2009; Zimmermann *et al*., 2020) might well complement its insulinotropic effect to further improve glycemic control.

The nature of sildenafil-induced Ca^2+^ influx is particularly intriguing, as it comprises a substantial amount of dihydropyridine (DHP)-insensitive components. In mouse β cells, glucose-induced Ca^2+^ influx is predominantly mediated by L-type VDCCs (Rorsman and Ashcroft, 2018; Thompson and Satin, 2021). Thus, treatment with DHP such as isradipine and nifedipine, or genetic ablation of CaV1.2, a subunit of L-type VDCC, profoundly suppresses GIIS (Schulla *et al*., 2003). However, there is also a DHP-insensitive component in GIIS, which is attributable to R-, P/Q-, and possibly N-type VDCCs in mouse β cells (Rorsman and Ashcroft, 2018; Thompson and Satin, 2021). Non-L-type VDCCs also modulate physiological insulin secretion, with R-type VDCCs participating in the second phase of GIIS (Jing *et al*., 2005). DHP-insensitive Ca^2+^ influx by sildenafil suggests that sildenafil directly activates non-L-type VDCCs or alternatively modulates the membrane potential to preferentially activate these channels. These characteristics may well be advantageous for use in human β cells, as DHP-insensitive Ca^2+^ currents appear to be more important for insulin secretion in humans than in mice (Davalli *et al*., 1996; Braun *et al*., 2008).

The known pharmacology of sildenafil involves increased intracellular cyclic guanosine monophosphate (cGMP) levels through inhibition of its hydrolysis via PDE5. Elevated cGMP levels activate protein kinase G (PKG), leading to various cellular responses. Indeed, research has suggested that cGMP/PKG activation can enhance insulin secretion through PKG-mediated K_ATP_ channel inhibition (Ropero *et al*., 1999) or membrane depolarization by unidentified K^+^ channels (Ishikawa *et al*., 2003). In addition, sildenafil can increase Ca^2+^i influx by cGMP/PKG-dependent activation of Ca^2+^-activated K^+^ channels with large conductance (BK channels) in human umbilical vein endothelial cells (HUVEC) (Luedders *et al*., 2006). However, these pathways do not seem to be involved in sildenafil-amplified insulin secretion, insofar as *Pde5a* knockdown, which might be expected to impede the ability of sildenafil to boost cGMP levels, in fact enhanced sildenafil responsiveness, indicating that sildenafil-amplified insulin secretion and cGMP/PKG signaling are not correlated.

This study has several limitations. First, the results are based solely on immortalized clonal β cells, which may not accurately represent the function of primary β cells in vivo. Further confirmation using primary islets or β cells is warranted. Second, this study did not confirm its findings in vivo. While previous studies failed to observe any change in plasma insulin or C-peptide levels after chronic sildenafil administration in diet-induced obese mice (Ayala *et al*., 2007; Johann *et al*., 2018) or prediabetic humans (Ramirez *et al*., 2015), this lack of effect may be attributed to the pharmacological profile of sildenafil. We demonstrated that the drug must be present at concentrations greater than 10 μM to stimulate insulin secretion, whereas its plasma concentration in vivo is typically less than 1 μM (approximately 500 ng/mL) after a single dose, as determined by pharmacokinetic analysis (Nichols *et al*., 2002; Alwhaibi *et al*., 2021). Therefore, to bridge the gap between in vitro and in vivo studies, the administration protocol needs to be optimized.

## 7. CONCLUSION

Our results indicate that sildenafil increases insulin secretion by enhancing Ca^2+^ influx via a mechanism independent of L-type VDCCs or PDE5. This study presents a new perspective on the metabolic advantages of sildenafil and provides insights into the molecular mechanism of insulin secretion.

## AUTHOR CONTRIBUTIONS

Conceptualization, N.M.; Methodology, N.M..; Investigation, N.M., R.M., K.S., and Y.M.; Writing – Original Draft: N.M. and R.M.; Writing – Review & Editing: N.M., R.M., K.S., Y.M., Y.S., Y.Y., and A.S.; Data Curation: N.M. and R.M.; Visualization: N.M. and R.M.; Supervision: K.S., Y.S., Y.Y., and A.S.; Funding Acquisition: N.M. and A.S.

## ACKNOWLEDGEMENTS

The authors extend their gratitude to President Yutaka Seino of Kansai Electric Power Hospital for his general support in this research. The authors thank Shihomi Sakai for her excellent technical assistance.

## DATA AVAILABILITY

Data supporting the findings of this study are available from the corresponding author upon reasonable request.

